# ADHD Medications and Preadolescent Brain Structure: Patterns of Cortical Attenuation from the ABCD Study

**DOI:** 10.1101/2025.11.13.687018

**Authors:** L. Nate Overholtzer, Katherine L. Bottenhorn, Sarah L. Karalunas, Bradley S. Peterson, Megan M. Herting

**Affiliations:** USC-Caltech MD-PhD Program, Keck School of Medicine of USC, Los Angeles, CA, USA; Neurosciences Graduate Program, University of Southern California, Los Angeles, CA, USA; Department of Population and Public Health Sciences, Keck School of Medicine of USC, Los Angeles, CA, USA; Department of Psychological Sciences, Purdue University, West Lafayette, IN, USA; Institute for the Developing Mind, Children’s Hospital Los Angeles, Los Angeles, CA, USA; Department of Psychiatry, Keck School of Medicine of USC, Los Angeles, CA, USA

**Keywords:** ADHD, amphetamine, methylphenidate, nonstimulants, MRI

## Abstract

Attention-deficit/hyperactivity disorder (ADHD) is the most common neurodevelopmental disorder in the U.S., and the stimulant and nonstimulant medications used to treat ADHD are among the most widely prescribed treatments in youth. Stimulants—including amphetamine-based (AMP) and methylphenidate-based (MPH) medications—act primarily on dopaminergic and noradrenergic systems, while nonstimulants (NS) more selectively target noradrenergic pathways. Although pharmacotherapy is the most clinically effective treatment, its neurostructural effects remain poorly understood. Leveraging the Adolescent Brain Cognitive Development• Study (ABCD Study®), we used a machine learning approach to identify neuroanatomical targets of medications, followed by linear mixed-effects modeling to estimate the effects of ADHD status and medication class (AMP, MPH, NS) on cortical thickness, surface area, and cortical and subcortical volumes. ADHD was not associated with statistically significant differences; however, a consistent pattern emerged in which AMP and MPH effects attenuated ADHD effects, suggesting that stimulant medications may attenuate ADHD-related cortical patterns. NS medications showed a similar, albeit weaker, effect pattern. Notably, AMP and/or MPH use was associated with significant effects in the right entorhinal cortex and the right banks of the superior temporal sulcus, potentially reflecting overcompensatory effects, as well as in the left posterior cingulate, possibly indicating *de novo* medication-related differences.

## Introduction

Attention-deficit hyperactivity disorder (ADHD) is the most prevalent neurodevelopmental disorder in the United States, with estimates of the biological prevalence ranging from 2 to 8%.^1–3^ Untreated, ADHD can substantially interfere with normal cognitive and functional pediatric development.^4^ Among all treatment methods, pharmacologic therapy is the first-line treatment approach and has the most substantial evidence for improving symptoms of inattention, impulsivity, and hyperactivity, while reducing negative impacts on academic, social, and occupational functioning.^5,6^ Accordingly, approximately 6% of U.S. children ages 3 to 17 years are treated with one or more medications for ADHD, making these medications some of the most common neuropsychiatric medications prescribed during childhood.^7,8^

ADHD medications have been hypothesized to “normalize” differences in brain structure and function. Broadly, medications include stimulant (e.g., amphetamine-based [AMP] and methylphenidate-based [MPH]) medications as the preferred first-line agents and nonstimulant [NS] medications (e.g., norepinephrine reuptake inhibitors and alpha-2-adrenergic agonists) as second-line agents if patients fail to respond to stimulants, experience intolerable side effects, or prefer nonstimulant options. Despite stimulants being the first-line treatment, 40% of children with ADHD display a preferential response to either AMP or MPH, while 30% of children fail to respond.^9–11^ The reasons behind variability in ADHD medication response remain unclear.^5,11^ Understanding how these medications affect brain phenotypes may help advance precision biomarkers for predicting treatment response.^12^

Stimulant medications exert their effects primarily by increasing synaptic levels of dopamine (DA) and norepinephrine (NE) through the inhibition of transporters (e.g., DAT and NET). However, AMP and MPH differ in their additional mechanisms of action: AMP also inhibits monoamine breakdown pathways (e.g., VMAT2 and MAO), while MPH displays weak agonism of serotonergic 5-HT1A and alpha-2-adrenergic receptors.^13^ Nonstimulant medications, including alpha-2-adrenergic agonists (e.g., clonidine, guanfacine) and norepinephrine reuptake inhibitors (e.g., atomoxetine, viloxazine), primarily modulate NE and lack the direct DA activity of stimulants.

Despite the widespread use of ADHD medications in the U.S. and worldwide, research on their effects on MRI brain phenotypes has been constrained by small, homogeneous samples that lack the statistical power to capture nuanced effects on brain structure and sample diversity to produce generalizable findings. Relatively few studies have examined structural MRI (sMRI) outcomes related to ADHD medication use, in contrast to the larger body of functional MRI literature.^14^ Among sMRI studies, stimulant-associated attenuation has been observed in frontal, cingulate, and parietal-occipital cortical regions,^15,16^ in basal ganglia and cerebellar subregions,^17–19^ and in the corpus callosum and global white matter.^20,21^ However, generalizability across these studies remains poor, with inconsistent and null findings.^14^

While meta-analyses have linked stimulant treatment in pediatric ADHD to attenuation in right hemisphere volumes of basal ganglia subregions (e.g., globus pallidus, putamen, lentiform),^22,23^ they have failed to identify replicable brain differences attributable to ADHD pathology itself, likely due to inter-study variability.^24^ However, the prefrontal cortex and basal ganglia are among the most frequently reported grey matter regions exhibiting ADHD-related differences in the literature.^25,26^ These findings align well with the roles of DA and NE in mesocortical and noradrenergic pathways that support prefrontal function and executive control, as well as mesolimbic pathways involved in emotion and reward processing.^27^ These dopaminergic pathways modulate the affective and cognitive loops of the cortico-striato-thalamo-cortical (CSTC) circuit, where dysregulation is hypothesized to contribute to the pathophysiology of ADHD and morphology in related brain regions.^28,29^

Given differences in neurotransmitter pathways and regional brain variation in transporter abundance, secondary receptor interactions, and catabolic enzymes, it is plausible that stimulants and nonstimulants may achieve similar improvements in ADHD symptoms via distinct neurostructural changes (**Figure 1**).^13,30,31^ Howeve, no studies have directly compared the neurostructural effects of different ADHD medication classes (i.e., AMP vs. MPH vs. NS), and few have examined the brain effects of any single class using sufficiently powered samples.^14^ Thus, our objective was first to use an explainable artificial intelligence (xAI) framework for a data-driven approach to identifying potentially medication-sensitive brain regions, and then to characterize the distinct neurostructural effects of AMP, MPH, and NS in relation to ADHD status in these regions. We leveraged a large and demographically diverse sample of preadolescents from the NIH-funded Adolescent Brain and Cognitive Development• Study (ABCD Study®). We hypothesized that stimulants (AMP and MPH) would be associated with attenuation of ADHD-related morphological differences, particularly in the ventromedial prefrontal cortex, cingulate cortex, and basal ganglia. In contrast, we expected nonstimulant effects to be more limited, reflecting primarily noradrenergic pathways and possibly affecting parietal regions.

**Figure 1.**
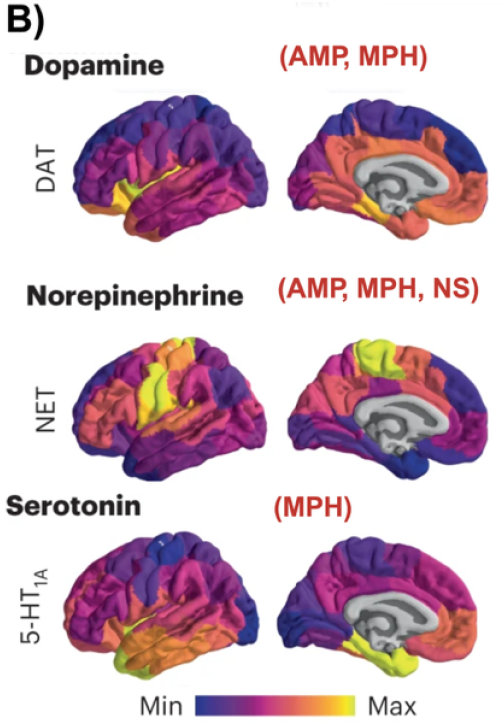
Schematic of neurotransmission systems relevant in ADHD pharmacology. **A)** Dopaminergic and noradrenergic pathways in the brain. Figure adapted from Faraone et. al, 2015.^30^ ***Note: Panel A of this figure is not displayed in the bioRxiv Preprint due to copyright laws*. B)** Neurotransmitter transporters and receptors relevant to ADHD medications exhibit varying concentrations throughout the cortex. Figure adapted from Hansen et al., 2022.^31^ **Abbreviations:** DA = dopamine (dopaminergic); NE = norepinephrine (noradrenergic); DAT= dopamine transporter; NET = norepinephrine transporter; 5-HT_1A_ = serotonin receptor subtype 1A; AMP = amphetamine; MPH = methylphenidate; NS = nonstimulant .

## Methods

### Participants

Cross-sectional data were collected as a part of the ongoing ABCD Study and included in the ABCD 5.1 data release (doi: 10.15154 / z563-zd24). The ABCD Study enrolled more than 11,800 children 9 and 10 years of age (mean age = 9.49; 48% female) between 2016 and 2018 for a 10-year longitudinal study.^32^ Participants were recruited at 21 study sites across the United States, aiming to represent nationwide sociodemographic diversity.^33^ Experimental and consent procedures are approved and overseen by the institutional review board (IRB) and human research protection programs at the University of California, San Diego. Each study site received local IRB approval. Participants provided written assent, and their legal guardian provided written consent to participate. See Garavan et al. (2018)^33^ and Volkow et al. (2018)^34^ for additional details. Criteria included English proficincy, absence of severe sensory, neurological, medical, or intellectual limitations, and completion of an MRI scan.^35^

The present study analyzes a subset of participant ABCD baseline enrollment data at ages 9 and 10 years, including structural magnetic resonance imaging (sMRI), ADHD diagnostic criteria, and medication usage. For this study, we excluded participants based on sMRI quality control procedures,^36^ incidental radiological findings,^37^ and other neurologic or psychiatric medications. Participants with missing covariate data were included in the machine learning (ML) stage of our analysis, which did not incorporate confounders; however, they were excluded from the LME modeling stage (N=5). Additionally, to address within-family correlation, we semi-randomly selected one sibling per family for the LME modeling stage, with the caveat that we preferentially retained the sibling with ADHD. Details of the present study’s exclusion criteria are in **Supplemental Figure 1**. Sample characteristics for our study are described in **Table 1**.

**Table 1.** Study Sample Characteristics.

|  | **Analysis 1:**<br>**ML-Sample**<br>**(N=1310)** | **Analysis 2:**<br>**LME-Sample**<br>**(N=8873)** |
| --- | --- | --- |
| **ADHD Classification** |  |  |
| ADHD+RX<sup>a</sup> | 753 (57.5%) | 699 (7.9%) |
| ADHD-NoRX<sup>b</sup> | 557 (42.5%) | 543 (6.1%) |
| Control | 0 (0.0%) | 7630 (86.0%) |
| **ADHD Medication Use** |  |  |
| AMP | 275 (21.0%) | 257 (2.9%) |
| MPH | 360 (27.5%) | 333 (3.8%) |
| NS | 161 (12.3%) | 151 (1.7%) |
| **Age (Months)** |  |  |
| Mean (SD) | 119 (7.43) | 119 (7.41) |
| Median [Min, Max] | 119 [107, 132] | 119 [107, 133] |
| **Sex at Birth** |  |  |
| Male | 916 (69.9%) | 4635 (52.2%) |
| Female | 394 (30.1%) | 4237 (47.8%) |
| **Race/Ethnicity** |  |  |
| Non-Hispanic white | < 680<sup>c</sup> | 4497 (50.7%) |
| Non-Hispanic Black | 227 (17.3%) | 1320 (14.9%) |
| Hispanic/Latinx | 244 (18.6%) | 1923 (21.7%) |
| Non-Hispanic Asian | < 10<sup>c</sup> | 204 (2.3%) |
| Other | 159 (12.1%) | 928 (10.5%) |
| **Household Income** |  |  |
| <\$50K USD | 420 (32.1%) | 2453 (27.6%) |
| \$50K to \$100K USD | 336 (25.6%) | 2287 (25.8%) |
| ≥\$100K USD | 447 (34.1%) | 3363 (37.9%) |
| Don't know/Refuse to answer | 107 (8.2%) | 769 (8.7%) |
| **Parent Education** |  |  |
| < HS Diploma | 56 (4.3%) | 460 (5.2%) |
| HS Diploma/GED | 135 (10.3%) | 880 (9.9%) |
| Some College | 410 (31.3%) | 2299 (25.9%) |
| Bachelor's degree | 331 (25.3%) | 2189 (24.7%) |
| Post Graduate Degree | 378 (28.9%) | 3034 (34.2%) |
| Missing/Refuse to Answer | 0 (0.0%) | 10 (0.1%) |
| **MRI Manufacturer** |  |  |
| GE Medical Systems | 303 (23.1%) | 2223 (25.1%) |
| Philips Medical Systems | 144 (11.0%) | 1103 (12.4%) |
| Siemens | 863 (65.9%) | 5546 (62.5%) |

<sup>a</sup> ADHD+RX = medicated ADHD group (i.e., participants using at least one ADHD medication)
<sup>b</sup> ADHD-noRX = unmedicated ADHD group (i.e., participants who met ADHD diagnostic criteria and did not report ADHD medication use)
<sup>c</sup> Small cell sizes (< 10) were masked in accordance with ABCD data-use guidelines. Secondary cell suppression is implemented to ensure category frequencies cannot be recalculated by subtraction. Percentages for masked or suppressed cells are not shown.
**Abbreviations:** ML = machine learning; LME = linear mixed-effects modeling; AMP = amphetamine; MPH = methylphenidate; NS = nonstimulant; HS = high school

#### ADHD and Medication Classification

Caregivers completed the computerized *Kiddie Schedule for Affective Disorders and Schizophrenia for DSM-5* (KSADS-5) to identify participants meeting ADHD diagnostic criteria, and the Medication History Questionnaire (MHQ), used to determine AMP, MPH, and/or NS medication use. ^38^ All participants taking antidepressant, antipsychotic, anxiolytic, antiepileptic, or other neurologic /psychiatric medications were excluded to reduce potential bias introduced by other neuroactive drugs or conditions. All participants currently using at least one ADHD medication were classified as the medicated ADHD group (ADHD+RX). This approach was necessary because KSADS does not account for whether reported symptoms reflect behavior while on or off medication, which could otherwise lead to a substantial under-classification of individuals with ADHD well-controlled by medication. From the ADHD+RX group, we identified and removed two subjects using an NS medication for Tic Disorder, rather than ADHD. Participants who met ADHD diagnostic criteria and did not report ADHD medication use were labeled as the unmedicated ADHD group (ADHD-noRX). The remaining participants were labeled the non-ADHD control (Control) group.

### Neuroimaging Data

Scanning protocol details have been reported by Casey et al. (2018).^39^ A harmonized MRI protocol was used across sites, utilizing 3T scanners manufactured by Siemens, Philips, or GE. Motion compliance training and real-time, prospective motion correction were used to minimize motion distortion. T1-weighted (T1w) images consisting of 176 slices with 1 mm^3^ isotropic resolution were acquired using a magnetization-prepared rapid acquisition gradient echo (MPRAGE) sequence.^39^ The ABCD Data Analysis, Informatics, and Resource Center inspected individual T1w images for poor image quality (*imgincl_t1w_include*) and incidence of abnormal clinical findings (*mrif_score*). The cortical surface of participants’ T1w images was reconstructed using FreeSurfer v7.1.1 and registered to the Desikan-Killaney Atlas to quantify cortical thickness, surface area, and volume across 68 cortical brain regions of interest (ROIs) and to the Aseg Atlas to quantify volumes in 14 subcortical ROIs.^36^

### Demographic Data and Covariates

The age, sex, and household size of the ABCD cohort closely match the distribution of 9- and 10-year-olds in the U.S. Census Bureau’s American Community Survey. The racial breakdown is similar, except for the underrepresentation of Asian, American Indian /Alaska Native, and Native Hawaiian /Pacific Islander children in the ABCD.^40^

Covariates for our study included these demographic and socioeconomic variables: child’s age (in *months*), child’s sex at birth (*male* or *female*), race /ethnicity (*Non-hispanic white, Non-hispanic Black, Hispanic/Latinx, Asian*, or *Other*), average household income (*••$100•K USD, ••$50•K and <•$100•K USD, <•$50•K USD*, or *Don’t Know/Refuse to Answer*), and highest household education (*Post-Graduate Degree, Bachelor Degree, Some College, High School Diploma/GED*, or *Less than High School Diploma*). The *Other* category of the combined race / ethnicity variable in ABCD includes caregiver-identified participants as American Indian /Native American, Alaska Native, Native Hawaiian, Guamanian, Samoan, other Pacific Islander, or other race.

### Analyses

Each analysis followed a two-step approach: (1) a region-of-interest (ROI) selection phase using bootstrap-enhanced elastic net (ENet) regression to identify regions potentially sensitive to medication effects (i.e., those that differ between medicated and unmedicated ADHD groups), and (2) linear mixed-effects (LME) regression modeling applied to the selected ROIs to estimate the effects of ADHD status and the specific medication classes (i.e., AMP, MPH, and NS). This workflow is depicted in **Figure 2** and was applied three times to measures of cortical thickness, surface area, and volume. All analyses were conducted in R Version 4.4.1 (R Core Team, 2024) using the *glmnet* version 4.1-8 package^41^ for ENet models and the *lme4* version 1.1-35.5 package^42^ for LME models, with model diagnostics assessed using the *performance* version 0.12.2 package.^43^

**Figure 2.**
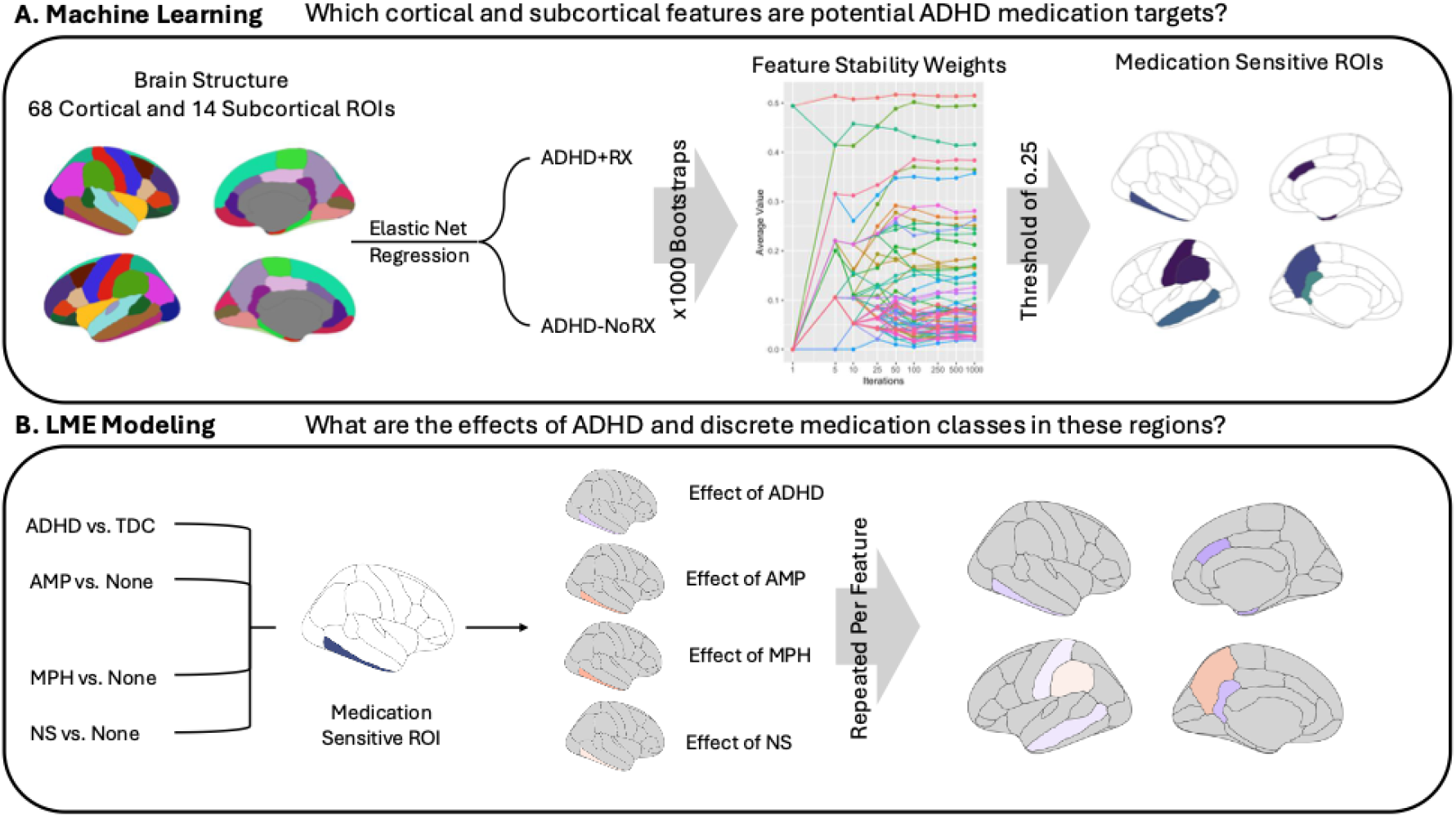
Workflow of our two-step analysis pipeline. **A)** First, a Machine Learning phase is used to identify brain regions that are potentially sensitive to ADHD medications. **B)** Second, LME models are used to measure the effects of ADHD and medication classes on identified regions. **Abbreviations:** ADHD+RX = medicated ADHD group; ADHD-noRX = unmedicated ADHD group; LME = mixed effects modeling; AMP = amphetamine; MPH = methylphenidate; NS = nonstimulant.

#### Analysis 1: Brain Feature Selection through Machine Learning

To identify potential neuroanatomic targets of ADHD medications, we applied bootstrap-enhanced elastic net (ENet) regression modeling to three sMRI metrics across 68 cortical ROIs and 14 subcortical ROI volumes to predict ADHD medication status (ADHD+RX vs. ADHD-noRX) as a logistic target. Each ML step was run separately for each sMRI metric (e.g., cortical thickness, surface area, volume). ENet regression was chosen because combining *L*_*1*_ and *L*_*2*_ regularization can accommodate the high degree of autocorrelation characteristic of MRI data. That is, it strikes a balance between sparsifying model coefficients (i.e., performing feature selection for dimensionality reduction) and retaining coefficients for correlated features (e.g., similar brain regions). This method, proposed by Bunea et al. (2011),^44^ involves repeatedly bootstrapping ENet regression and has been shown to improve MRI feature selection compared to traditional ENet regression.

Analyses were conducted in R (v4.4.1) using the *glmnet* (v4.1-8) package, which standardizes features. Specifically, we performed 1,000 bootstraps of cross-validated ENet regression, ensuring each iteration preserved a stratified ratio of ADHD+RX to ADHD-noRX subjects in both the 80% training and 20% testing sets. The *L*_*1*_:*L*_*2*_ ratio, or *•*, was set at 0.5 to ensure models balanced sparsity and shrinkage, as noted above. Within each iteration, *•* was fine-tuned via 10-fold cross-validation across the 80% training set, selecting the regularization parameter that yielded the lowest cross-validated mean squared error. Non-zero coefficients from each model iteration indicated whether an ROI was selected. To account for model quality, we weighted each selected ROI by its model’s performance on the held-out 20% testing set, using the area under the curve (AUC_Test_). These weighted selection scores were then averaged across bootstrap iterations to generate a feature stability weight for each ROI. Building on the approach of Bunea et al. (2011)^44^ and Abram et al. (2016)^45^ in retaining ROIs selected in 50% of bootstrap iterations, we set our feature stability threshold at 0.25—equivalent to selection in 50% of iterations weighted by a model performing at chance level (AUC_Test_ = 0.50).

#### Analysis 2: Linear mixed-effects (LME) modelling

Selected ROIs were included in the regression modeling stage to assess the effects of our primary dependent variables: (1) ADHD status, as well as medication class effects of (2) AMP, (3) MPH, and (4) NS on the brain in these ROIs (i.e., independent variables). ADHD status and each medication class were included as binary variables since some subjects display polypharmacy (i.e., use of more than one ADHD medication class). Each regression model included fixed effects of demographic covariates and MRI scanner manufacturer, with a random effect of MRI scanner serial number. Due to the small categorical counts, we combined the *Asian* and *Other* race / ethnicity categories into a single categorical variable (*Asian/Other*) for regression analysis. Inclusion of MRI scanner manufacturer (*Siemens, Philips*, or *GE*) accounts for differences in scanner hardware and software in analyses. Instead of an ABCD site effect, we included a random effect that accounted for the 29 unique MRI scanners using their serial numbers, which has been demonstrated to adequately model the significant scanner-related effects observed in the ABCD Study.^46^ See **Supplemental Methods** for model details. Additionally, we included total intracranial volume as a confounder in analyses of cortical and subcortical volumes. After removing participants with missing covariate data, we randomly selected one child per family. Importantly, there was no evidence of problematic collinearity for LME models, with Variance Inflation Factor (VIF) values of 2.01 for ADHD, 1.34 for AMP, 1.50 for MPH, and 1.17 for NS in our data. False discovery rate (FDR) correction was performed separately for each predictor (e.g., ADHD, AMP, MPH, NS) across each set of models (i.e., cortical thickness, surface area, volume) using the Benjamini-Hochberg method.^47^ Effect sizes are reported as standardized beta coefficients, allowing for direct comparison across brain features. Given the limitations of relying solely on statistical significance in large population-based studies,^48,49^ we first describe patterns in the beta coefficients—comparing the direction and average magnitude of effects for ADHD status with those of AMP, MPH, and NS—before reporting statistical significance.

## Results

### Sample Characteristics

Consistent with prior epidemiological findings in ADHD,^7^ our sample included a higher prevalence of male than female youth in both the medicated (Male: 73.1%; Female: 26.9%) and unmedicated ADHD groups (Male: 66.9%; Female: 32.9%), along with fewer participants identifying as Asian. Additionally, in line with prior findings,^7^ females and Hispanic /Latinx individuals within the ADHD groups were less likely to receive ADHD medication treatment. See **Supplemental Table 1**.

#### Analysis 1: Brain Feature Selection through Machine Learning

### Cortical and Subcortical Brain Regions Relevant to ADHD Medication Status

Under our feature stability weight threshold of 0.25, our ML approach identified nine regions for cortical thickness, two regions for cortical surface area, and twelve regions for cortical and subcortical volumes relevant for differentiating medicated and unmedicated ADHD participants (**Figure 3)**. Regions exhibiting relevant cortical thickness differences between ADHD+RX and ADHD-noRX individuals included the right caudal anterior cingulate, right entorhinal, right inferior temporal, left isthmus cingulate, left middle temporal, left precuneus, left postcentral, left supramarginal, and right temporal pole. The average AUC_Test_ across bootstrap iterations was 0.51 for cortical thickness. For surface area, relevant regions included the right banks of the superior temporal sulcus and the left posterior cingulate; the average AUC_Test_ was 0.54. For cortical and subcortical volume, relevant regions included the right banks of the superior temporal sulcus, right cuneus, left inferior parietal, right inferior temporal, right lingual, left pars triangularis, left posterior cingulate, left superior parietal, right temporal pole, right accumbens area, left amygdala, and right putamen; the average AUC_Test_ was 0.51. The average AUCs for our three bootstrap-enhanced ENet models were all near chance, underscoring the limitations of standard ENet models in reliably identifying features. This highlights the importance of bootstrap-enhancement, which improves feature importance metrics (e.g., our feature stability weight) by measuring feature selection frequency across resampled datasets. Post hoc, we discovered that approximately 100 bootsraps would have been sufficient for our data, as the feature stability weight leveled off around this point (**Supplemental Figures 2-4**).

**Figure 3.**
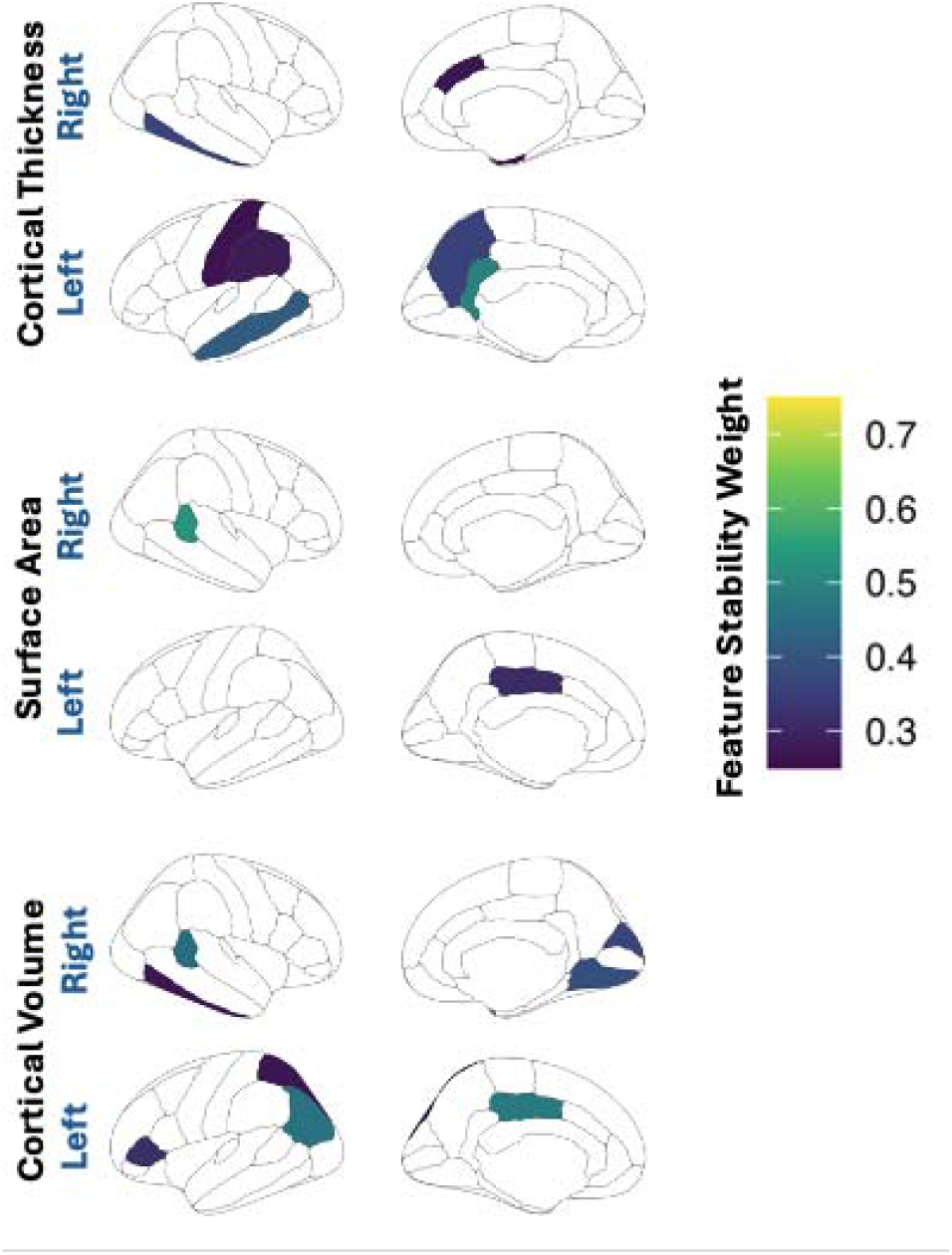
Brain Features Relevant to ADHD Medication Usage. Feature Stability Weights of brain regions predicting ADHD medication status (i.e., ADHD+RX vs. ADHD–NoRX) using bootstrap-like ENet regression. A feature stability weight of 0.25 was used as our threshold; a higher feature stability weight indicates a more reliable selection across bootstraps and better model performance. **Abbreviations:** ADHD+RX = medicated ADHD group; ADHD-noRX = unmedicated ADHD group.

#### Analysis 2: Linear mixed-effects (LME) modelling

Under our data-driven ML approach, brain regions identified as potentially medication-sensitive were then used in linear mixed-effects (LME) modeling to measure the effects of (1) ADHD Status, (2) AMP use, (3) MPH use, and (4) NS use. To explore broad, non-inferential patterns of ADHD and medication effects on brain structure, we describe the direction and magnitude of standardized beta coefficients across regions before considering statistical significance.

### Patterns in ADHD and Medications Effects on Brain Structure

All standardized beta coefficients for the primary predictors—regardless of statistical significance—are mapped across brain regions for cortical thickness, surface area, and volume in **Figure 4**, and for subcortical volume in **Supplemental Figure 5** to illustrate the broader patterns of associations between ADHD and medication effects with brain structure outcomes. These effects are also plotted with 95% confidence intervals in **Figure 5**.

**Figure 4.**
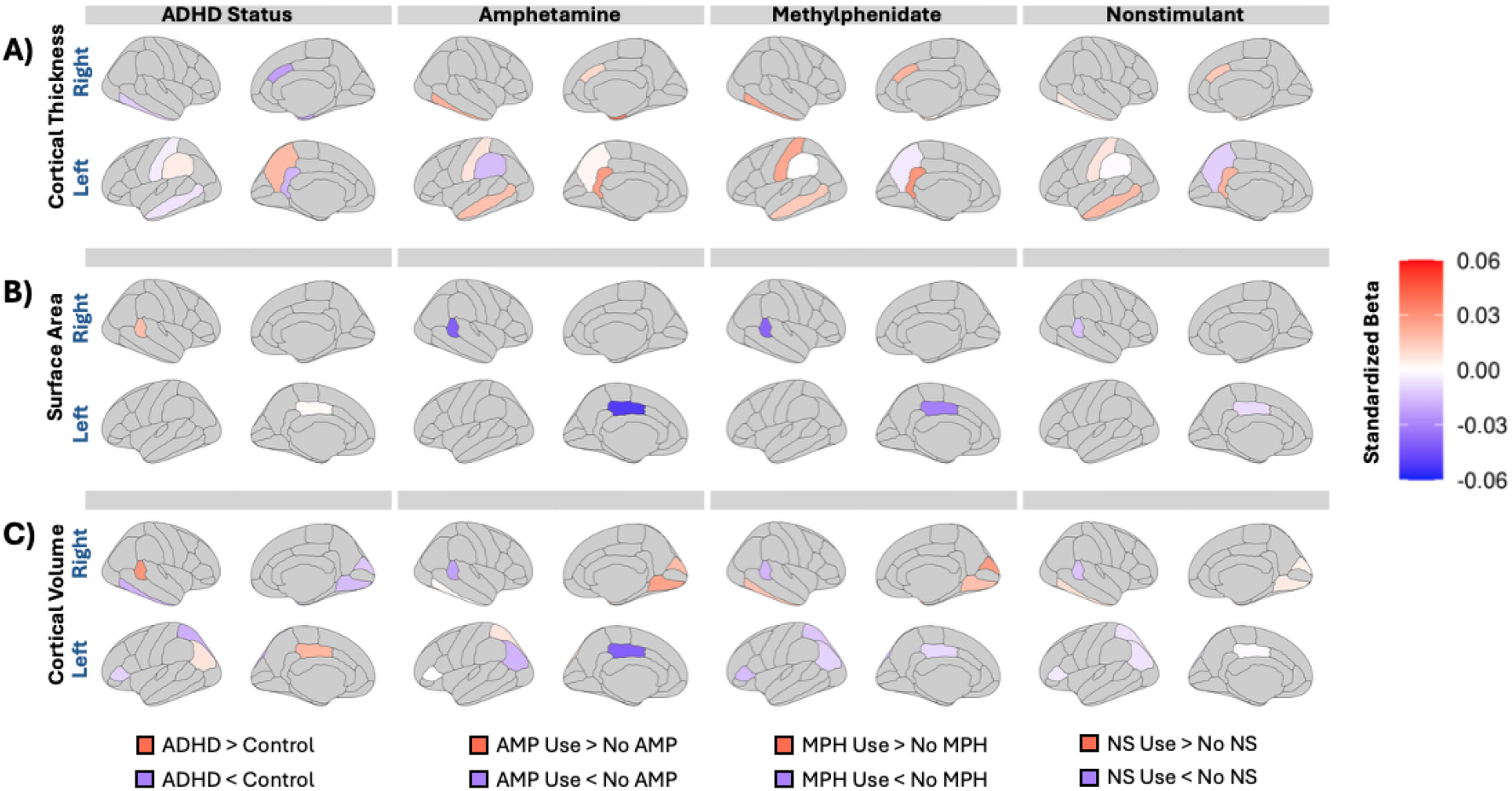
Effects of ADHD and medications mapped onto the Desikan-Killiany Atlas. Standardized beta coefficients for ADHD status, amphetamine (AMP), methylphenidate (MPH), and nonstimulant (NS) use from LME regression models across cortical brain features selected by ML, including a) cortical thickness, b) surface area, and c) cortical volume. Red indicates positive beta coefficients (i.e., higher values in the ADHD or medication-exposed groups), and blue indicates negative coefficients. Cortical regions not selected during the ML stage (and therefore not included in LME modeling) are colored in grey. **Abbreviations:** AMP = amphetamine; MPH = methylphenidate; NS = nonstimulant.

**Figure 5.**
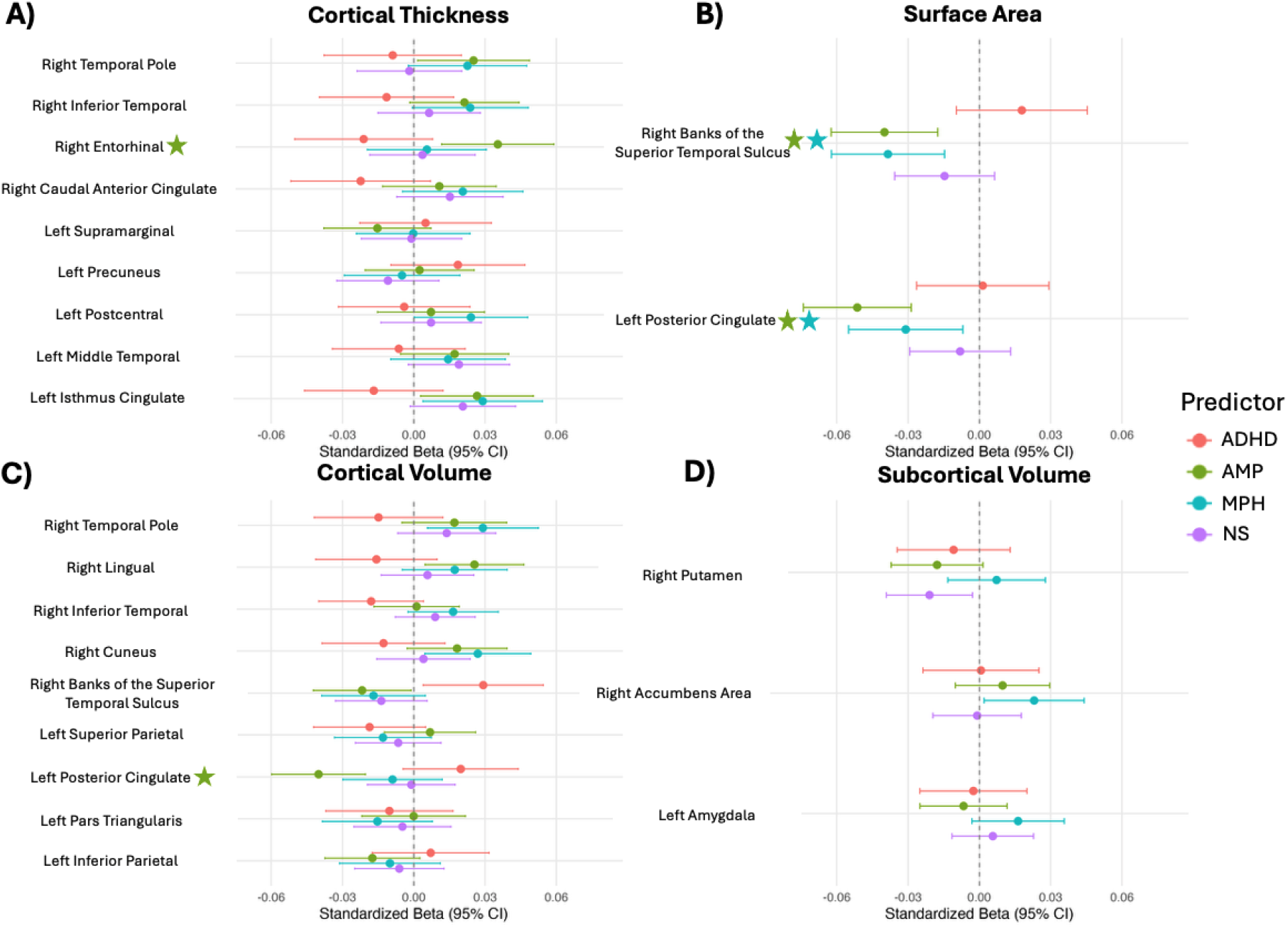
Forest plots of the effects of ADHD and medications. Standardized beta coefficient with 95% confidence intervals plotted for ADHD status, amphetamine (AMP), methylphenidate (MPH), and nonstimulant (NS) use from LME regression models for ML-selected cortical a) thickness, b) surface area, c) volume, and d) subcortical volume brain features. Effects that reached statistical significance after FDR correction are denoted with a star. **Abbreviations:** AMP = amphetamine; MPH = methylphenidate; NS = nonstimulant.

For cortical thickness, individuals with ADHD showed smaller cortical thickness compared to non-ADHD controls in the right caudal anterior cingulate, right inferior temporal, left isthmus cingulate, left middle temporal, left postcentral, and right temporal pole (**Figure 4A** and **Figure 5A**). In contrast, individuals with ADHD showed greater cortical thickness compared to controls in the right entorhinal cortex, left precuneus, and left supramarginal gyrus. AMP use was related to attenuation of ADHD cortical thickness differences (i.e., opposite directionality of ADHD status effect; or in other words, improvement compared to ADHD phenotype) in 8 of the 9 ROIs, with the exception of the left precuneus. Similarly, MPH use was related to attenuation in all 9 of the ROIs. NS use was also related to attenuation in 8 of the 9 ROIs, with the exception of the right temporal pole **(Figure 4A** and **Figure 5A**). Across the 9 cortical thickness ROIs, the average absolute values of the standardized betas were 0.013 ± 0.007 for ADHD, 0.018 ± 0.010 for AMP, 0.016 ± 0.010 for MPH, and 0.0095 for NS.

For surface area, individuals with ADHD had a larger surface area of the right banks of the superior temporal sulcus and left posterior cingulate (**Figure 4B** and **Figure 5B**). AMP, MPH, and NS use attenuated these differences in both ROIs. For surface area ROIs, the average absolute values of the standardized beta coefficients were 0.010 ± 0.012 for ADHD, 0.046 ± 0.008 for AMP, 0.035 ± 0.005 for MPH, and 0.010 ± 0.007 for NS.

For cortical and subcortical volumes, individuals with ADHD had smaller volumes in the right cuneus, right inferior temporal, right lingual, left pars triangularis, left superior parietal, right temporal pole, left amygdala, and right putamen (**Figure 4C, Figure 5C-D, Supplemental Figure 5**). In addition, those with ADHD had larger volumes as compared to controls in the right banks of the superior temporal sulcus, left inferior parietal, left posterior cingulate, and right accumbens area. AMP use attenuated ADHD effects in 8 of 12 ROIs, MPH use in 9 of 12 ROIs, and NS use in 9 of 12 ROIs. The primary exception for medication effects was the volume of the left pars triangularis, which was not attenuated by any of the three medication classes. Medications showed mixed associations with volumes in the left superior parietal cortex, right accumbens area, left amygdala, and right putamen, with at least one medication in the attenuating direction and at least one in the ADHD direction. Not considering the three subcortical regions, the medication attenuation rate for the cortical volumes was like that of cortical thickness and surface area. Across these cortical and subcortical volume ROIs, the average absolute standardized beta coefficients were 0.013 ± 0.008 for ADHD, 0.016 ± 0.011 for AMP, 0.017 ± 0.007 for MPH, and 0.008 ± 0.006 for NS.

### Significant Stimulant Effects in Cortical Regions

After false discovery rate (FDR) correction, significant effects were observed only for AMP and MPH use in cortical regions (denoted with a star in **Figure 5)**. Both stimulants were associated with reduced surface area in the right banks of the superior temporal sulcus and the left posterior cingulate (**Figure 5B**). Additionally, AMP use was linked to increased cortical thickness in the right entorhinal cortex and decreased volume in the left posterior cingulate following FDR correction (**Figure 5A** and **5C**). No effects of ADHD status or NS use reached significance after FDR correction. Additionally, no significant effects on subcortical volumes were observed after FDR correction.

## Discussion

This study is among the first to investigate brain structure differences associated with ADHD medication use in a large, demographically diverse sample. By integrating machine learning to identify medication-sensitive regions with linear mixed-effects regression modeling, we demonstrate opposing effects of ADHD status and medication use in parietal, occipital, and temporal regions. Our analytic approach is further strengthened by a theoretically grounded framework that allows us to disentangle the distinct effects of different ADHD medication classes while accounting for the common polypharmacy (e.g., AMP+NS or MPH+NS) observed in real-world populations.

### Medications Appear to Attenuate ADHD Brain Differences

We highlight the utility of our conceptual approach, which utilizes machine learning (ML) to identify potential medication-sensitive regions in youth with ADHD, within an explainable artificial intelligence (xAI) framework. We then further investigated how these subjects differ from those without ADHD (controls) and the effects of specific medication classes on morphology. The resulting attenuation patterns are strengthened by the fact that the ML phase was conducted without data from controls, and the selected brain features were identified solely for their ability to distinguish medicated from unmedicated individuals with ADHD. Notably, the morphology of many medication-sensitive brain regions appeared to be normalized (i.e., more like that of controls) by medication use, highlighting the utility of our combined data- and theory-driven approach. The cortical regions identified as likely medication targets were primarily located within the parietal, occipital, and temporal lobes, with a relative paucity of involvement in the frontal lobe, despite long-standing hypotheses about the role of prefrontal regions in ADHD pathology.^25,26^ One possibility is that prefrontal regions continue to exhibit ADHD-related structural differences but are not the primary targets of pharmacological intervention, that these differences in prefrontal regions emerge later during adolescence,^50^ or that their extended developmental trajectory into early adulthood^51^ may render them less susceptible to medication effects during the period of preadolescence studied here.

In our study of preadolescents, we found no statistically significant effects attributable to ADHD across the cortical and subcortical features examined when accounting for medication use, consistent with prior meta-analytic findings.^24^ Rather than relying solely on significance, we also examined the magnitude and direction of brain morphology associated with ADHD. In doing so, we observed both larger and smaller cortical brain region features relative to controls, highlighting a lack of global effects of ADHD at this age. Furthermore, when focusing on the patterns regarding the direction of effects, our findings align with prior research suggesting that ADHD medications may help “normalize” cortical structure.^14^ Moreover, AMP and MPH exhibited parallel patterns across cortical regions, with the direction of the effects aligning with potential attenuation of ADHD-related structural differences when compared to controls. For instance, although participants with ADHD showed reduced cortical thickness in several regions of the temporal lobe and the cingulate cortex when compared to control youth, stimulant usage was associated with relative increases in cortical thickness in these regions. NSs showed a similar pattern of attenuation, but with smaller beta coefficients compared to stimulants, suggesting weaker effects. However, this attenuation pattern for ADHD medications was not observed in subcortical regions, where medication effects appeared more variable.

Prior ABCD Study research by Wu et al. (2024)^52^ identified differences in right insula thickness and left nucleus accumbens volume between stimulant-treated individuals with low ADHD symptoms (N = 273) and unmedicated individuals with ADHD. However, these regions were not identified as potential primary medication targets in our study, potentially due to differences in analytic approach or the broader scope of our investigation—such as disaggregating stimulant classes, including a larger number of stimulant users, and examining both stimulant and nonstimulant medication effects. Additionally, in a small ABCD Study subsample, Kaminski et al. (2024)^53^ reported that stimulant use was associated with altered functional connectivity between striatal regions and several large-scale brain networks, including the cingulo-opercular, salience, fronto-parietal, cingulo-parietal, somatomotor, dorsal attention, ventral attention, and visual networks. While our ML approach identified two potential striatal targets of ADHD medications (the right putamen and right accumbens area), we found no significant effects of ADHD or medication use in these regions after FDR correction. Moreover, we observed no consistent patterns of ADHD attenuation between medication classes in subcortical regions. Despite the basal ganglia’s frequent implication in ADHD pathogenesis,^25,26^ our findings suggest that these medications may not alter striatal region structure during preadolescence, or that such effects may not lead to such morphological differences as assessed here.

### Stimulants May Overcompensate

Across medication-sensitive brain regions, the average absolute effects of AMP and MPH on cortical thickness, surface area, cortical volume, and subcortical volume were larger than those associated with ADHD. Interestingly, several cortical regions—such as the right entorhinal cortex, the right banks of the superior temporal sulcus, and the left posterior cingulate—showed statistically significant effects of AMP and /or MPH. In the entorhinal cortex and banks of the superior temporal sulcus, these effects were in the direction of attenuation, suggesting a potential normalization of ADHD-related structural differences. However, in these regions, the magnitude of stimulant-related effects exceeded those associated with ADHD diagnosis, raising the possibility of neurobiological overcompensation. NS medications exhibited similar directional effects to stimulants but with smaller magnitudes, consistent with their distinct pharmacological profile—namely, their primary action on noradrenergic pathways and downstream engagement of dopaminergic pathways.^54^ This also aligns with clinical evidence indicating that NSs generally yield more modest improvements in ADHD symptoms.^5^

Bernanke et al. (2022),^55^ using data from the same wave of the ABCD Study, identified reduced posterior cingulate surface area as the largest ADHD-related structural difference, but they did not account for ADHD medication use. In contrast, our study found significant reductions in left posterior cingulate surface area associated with both amphetamine (AMP) and methylphenidate (MPH) use, while the effect of ADHD in this region was near zero. This suggests that stimulants may relate to *de novo* alterations in the posterior cingulate, raising the possibility that previously reported differences may reflect medication effects rather than intrinsic ADHD-related pathology. Additionally, we observed that AMP use was associated with reduced posterior cingulate volume, implying that volumetric normalization may be primarily driven by surface area reduction rather than cortical thinning. Given that total surface area peaks during preadolescence, this morphometric feature may represent a particularly sensitive target for pharmacological effects during this developmental period.^51^

AMP exhibited more statistically significant effects, including unique associations with entorhinal cortical thickness and posterior cingulate volume, but not significant with MPH. This may reflect pharmacodynamic differences, as some literature suggests AMP leads to greater cytosolic dopamine accumulation than MPH.^56^ Because the spatial distribution of AMP-only effects (e.g., entorhinal cortex thickness, posterior cingulate volume) aligns with regions of higher dopamine transporter (DAT) density, these observed stimulant effects may be driven by dopaminergic mechanisms.^31^ In contrast, stimulant effects observed in the superior temporal sulcus may be driven more by noradrenergic mechanisms, given its higher relative expression of norepinephrine transporter (NET).^31^ Altogether, these findings emphasize the need to account for medications as key confounding variables in future ADHD neuroimaging research.

Our flexible LME modeling approach enabled us to isolate the unique effects of ADHD, AMP, MPH, and NS medications while handling polypharmacy. Unlike conventional approaches (e.g., grouping participants into mutually exclusive medication categories, aggregating stimulants, or excluding individuals with ADHD polypharmacy), our framework accommodates overlapping medication use. This approach reduces collinearity, preserves sample size, and more accurately reflects real-world treatment patterns, thereby enhancing interpretability and ecological validity. However, in theory, the observed medication-related effects on brain structure could also reflect unmeasured confounders related to medication treatment. For example, these findings could stem from who initiates treatment (e.g., individuals with brain features linked to greater severity or specific symptom profiles) or who maintains treatment (e.g., those with features associated with stimulant responsivity). Future longitudinal research is essential to determine whether these brain structure differences precede or follow the initiation of stimulant use.

### Strengths and Limitations

Although it lacks the rigor of a randomized controlled trial, the large and demographically diverse ABCD Study cohort offers an unparalleled opportunity to investigate pharmacologic effects under naturalistic conditions, including imperfect medication adherence. Additionally, our xAI framework used a data-driven feature selection approach to identify potential neuroanatomical targets of medications. This focus of our xAI framework may limit the identification of regions relevant to ADHD that are relatively independent of medication effects. Finally, to our knowledge, this is the first structural MRI study to model major ADHD medication classes concurrently by disaggregating stimulant classes and including nonstimulant medication effects. This has enabled us to identify both common and distinct neuroanatomical effects of these medications, representing a critical advancement in precision psychiatric medicine.

A key limitation of our analysis is its focus on a single developmental time point (ages 9–10 years) within a longitudinal study. ADHD has been hypothesized to reflect brain developmental delays, with phenotypic differences becoming more pronounced over the course of adolescence. Future longitudinal work is needed to assess how ADHD- and medication-related neurobiological differences evolve or resolve across later developmental stages. While our classification approach captures all individuals prescribed ADHD medication, it remains possible that some cases reflect overdiagnosis within the community. The ABCD Study does not survey medication history prior to the study period; therefore, the lifetime exposure to ADHD medications is unknown. We did not examine the potential moderating effects of medication dosage or disease severity, which are priorities for future research. Finally, due to the relatively small NS subsample size, we analyzed NS effects as a single categorical class, rather than disaggregating alpha-agonists and norepinephrine reuptake inhibitors, which are pharmacologically distinct.

## Conclusions

Our study employed an innovative approach to mapping the effects of ADHD medications on preadolescent cortical and subcortical brain development, identifying novel medication-related associations. We contribute several key findings to the field: (1) AMP and MPH appear to broadly attenuate ADHD-related effects in the cortex, and (2) NS exhibit similar but weaker effects to those of stimulants. Additional longitudinal research is needed to investigate whether stimulants may be associated with overcompensatory effects in specific regions of the cortex, particularly in the posterior cingulate, banks of the superior temporal sulcus, and entorhinal cortex, which may be due to their roles in dopaminergic and noradrenergic pathways.

## Acknowledgments

A special thanks to the participants and families of the ABCD Study. We would like to acknowledge the ABCD Consortium staff for their efforts in collecting data. We would also like to acknowledge Alethea de Jesus and Yara Akiel, who contributed to the data cleaning and formatting of the ABCD tabulated data prior to data analysis.

## Author Contributions

Conceptualization: LNO, KLB, SLK, BSP, MMH. Data Curation: LNO.

Formal Analysis: LNO.

Funding Acquisition: MMH.

Investigation: LNO.

Methodology: LNO, KLB.

Supervision: MMH.

Visualization: LNO.

Writing – Original Draft Preparation: LNO, MMH. Writing – Review & Editing: LNO, KLB, SLK, BSP, MMH.

## Data Availability Statement

Data used in the preparation of this article were obtained from the Adolescent Brain Cognitive Development (ABCD) Study (https://abcdstudy.org), held in the NIMH Data Archive (NDA). This is a multisite, longitudinal study designed to recruit more than 10,000 children aged 9–10 and follow them over 10 years into early adulthood. The ABCD Study is supported by the National Institutes of Health Grants [U01DA041022, U01DA041028, U01DA041048, U01DA041089, U01DA041106, U01DA041117, U01DA041120, U01DA041134, U01DA041148, U01DA041156, U01DA041174, U24DA041123, U24DA041147]. A complete list of supporters is available at https://abcdstudy.org/nih-collaborators. A list of participating sites and a comprehensive list of study investigators can be found at https://abcdstudy.org/principal-investigators.html. ABCD consortium investigators designed and implemented the study and/or provided data, but did not necessarily participate in the analysis or writing of this report. This manuscript reflects the views of the authors and may not reflect the opinions or views of the NIH or ABCD consortium investigators. The ABCD data repository grows and changes over time. Qualified researchers can request access to ABCD shared data from the ABCD Data Access Committee (DAC).

