## Supplement for "ADHD Medications and Preadolescent Brain Structure: Patterns of Cortical Attenuation from the ABCD Study"

This document includes:

- Supplemental Methods
- Supplemental Figure 1: Classification and Exclusion Flowchart
- Supplemental Figure 2: Cortical Thickness Average Feature Stability Weight by Iteration
- Supplemental Figure 3: Surface Area Average Feature Stability Weight by Iteration
- Supplemental Figure 4: Cortical and Subcortical Volume Average Feature Stability Weight by Iteration
- Supplemental Figure 5: Effects of ADHD and medications mapped onto the Aseg Atlas
- Supplemental Table 1: Descriptive Statistics of the ABCD Study Population by ADHD Classification

### Supplemental Methods

*Statistical Model Equations:*

Eq 1. Cortical Thickness:

Cortical Thickness of ROI ~ $ADHD_{Yes/No}$ + $AMP_{Yes/No}$ + $MPH_{Yes/No}$ + $NS_{Yes/No}$ + Sex + Age + Race/Ethnicity + Household Income + Parent Education + MRI Manufacturer + (1|MRI Serial No.)

Eq 2. Surface Area:

Surface Area of ROI ~ $ADHD_{Yes/No}$ + $AMP_{Yes/No}$ + $MPH_{Yes/No}$ + $NS_{Yes/No}$ + Sex + Age + Race/Ethnicity + Household Income + Parent Education + MRI Manufacturer + (1|MRI Serial No.)

Eq 3. Cortical and Subcortical Volumes:

Cortical/Subcortical Volume of ROI ~ $ADHD_{Yes/No}$ + $AMP_{Yes/No}$ + $MPH_{Yes/No}$ + $NS_{Yes/No}$ + Sex + Age + Race/Ethnicity + Household Income + Parent Education + MRI Manufacturer + ICV + (1|MRI Serial No.)

**Supplemental Figure 1.** Flowchart of subject selection detailing exclusion criteria applied to result in our Machine Learning (ML) sample and Linear Mixed-Effect modeling (LME) sample.

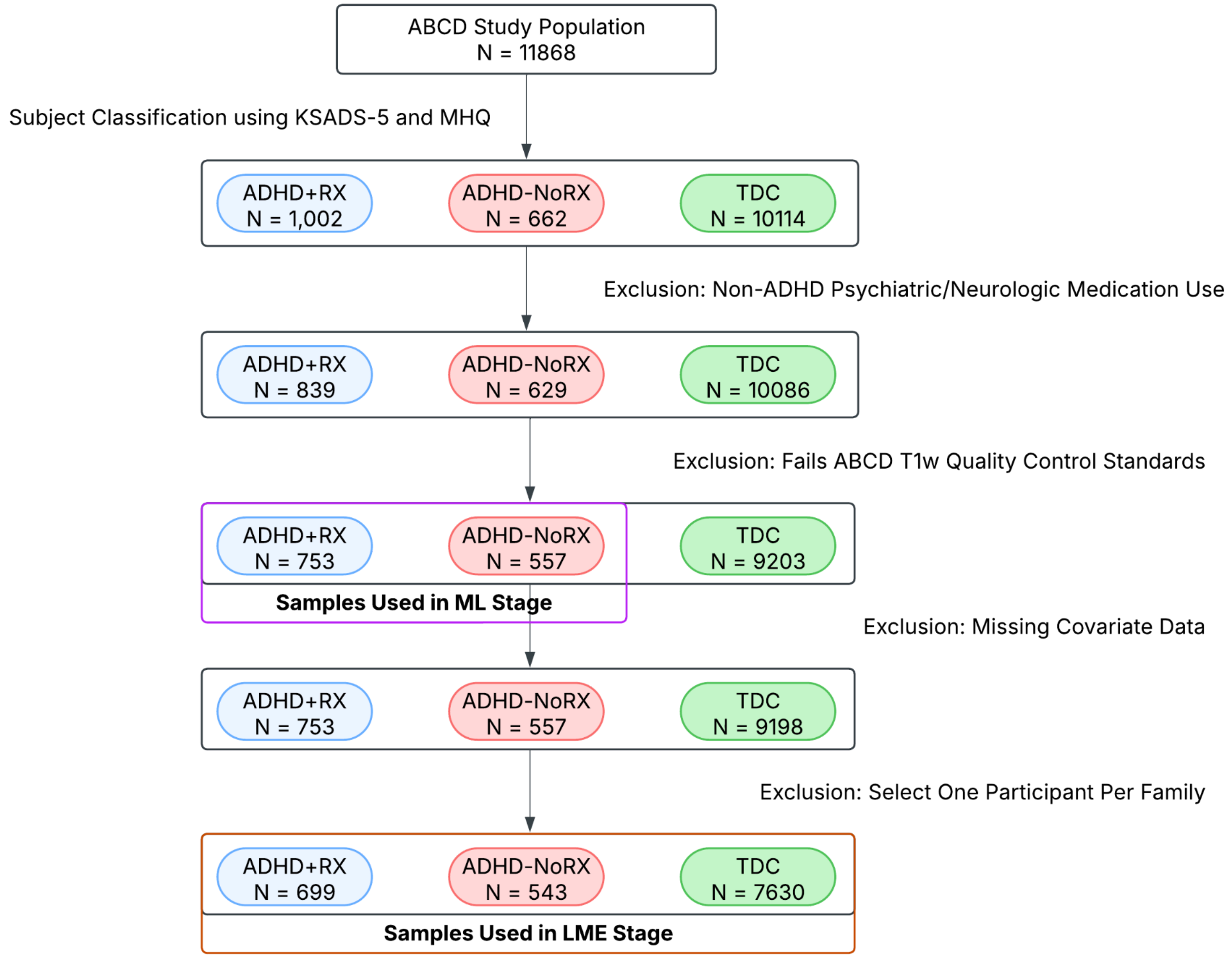

**Supplemental Figure 2.** Average Feature Stability Weight over 1000 Bootstrap Iterations for Cortical Thickness ROIs.

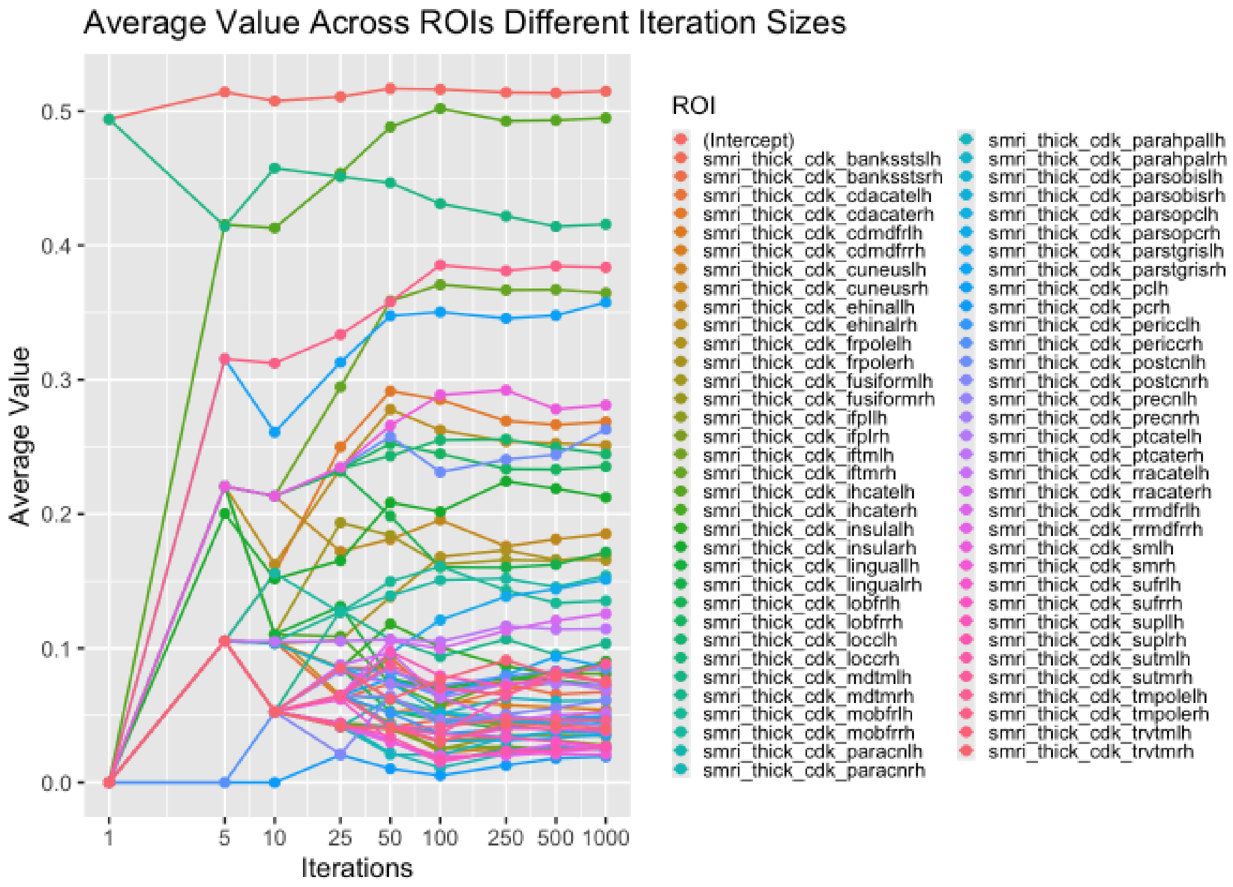

**Supplemental Figure 3.** Average Feature Stability Weight over 1000 Bootstrap Iterations for Surface Area ROIs.

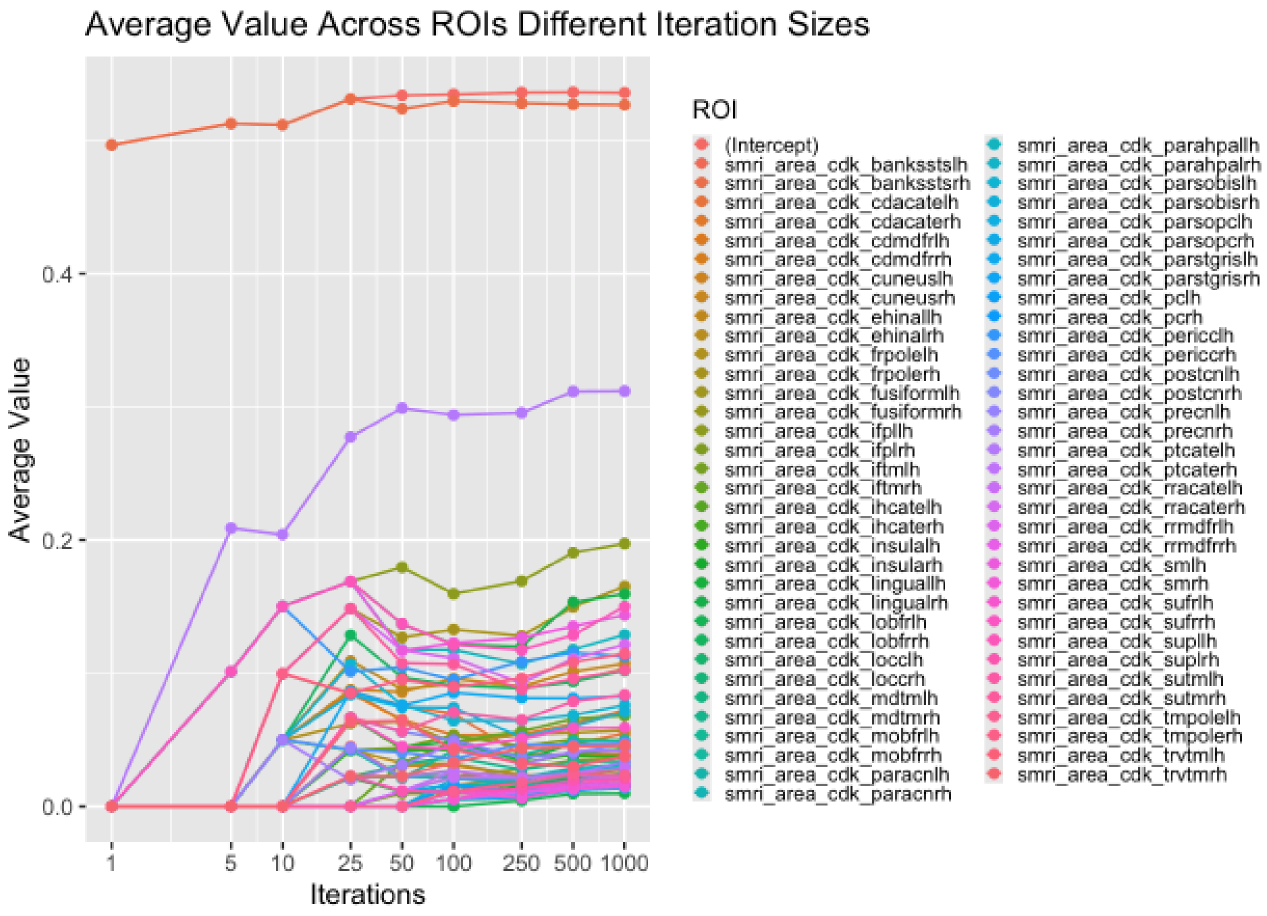

**Supplemental Figure 4.** Average Feature Stability Weight over 1000 Bootstrap Iterations for Cortical and Subcortical Volume ROIs.

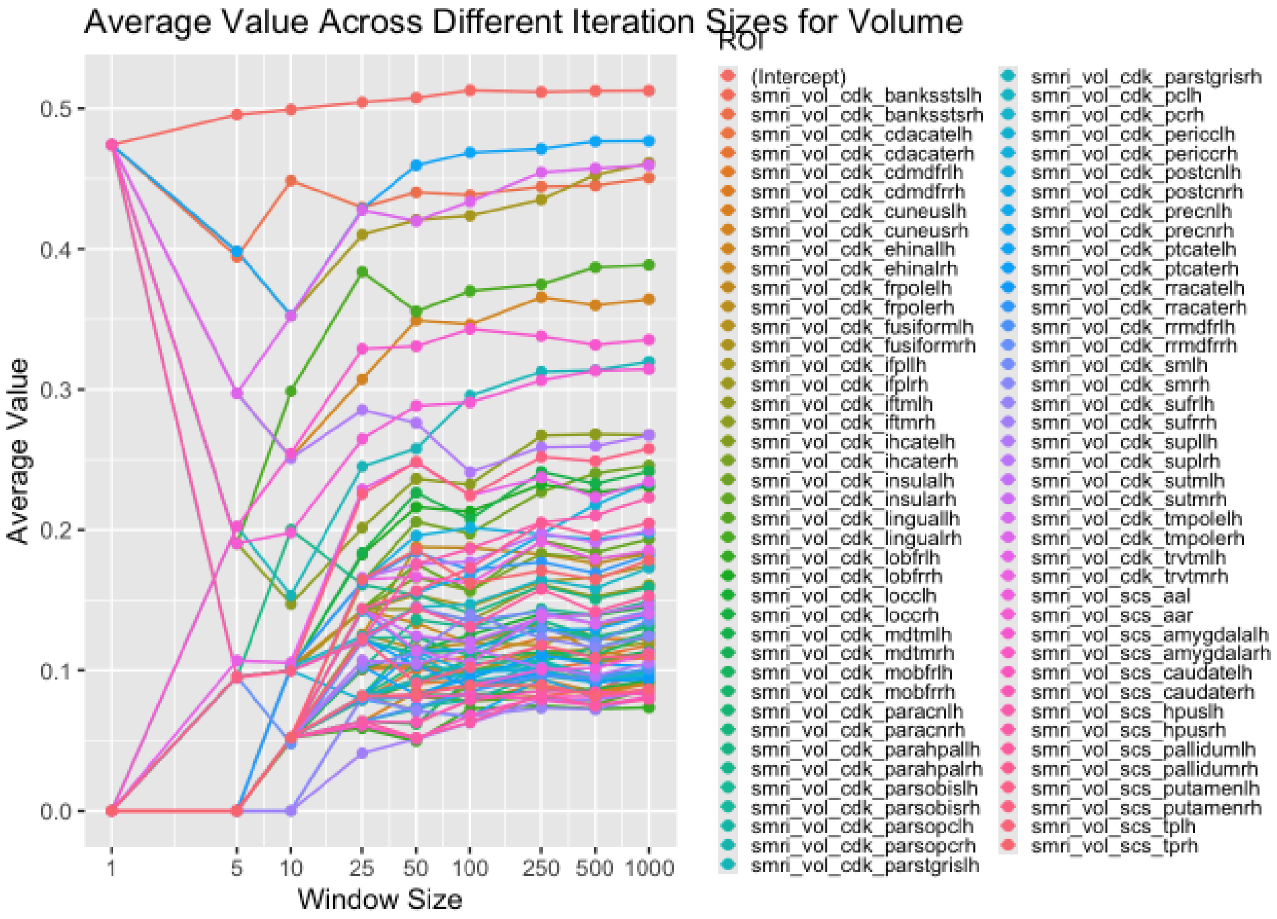

**Supplemental Figure 5. Effects of ADHD and medications mapped onto the Aseg Atlas.** Standardized beta coefficients for ADHD status, amphetamine (AMP), methylphenidate (MPH), and nonstimulant (NS) use from regression models across subcortical brain features selected during the machine learning (ML) stage. Red indicates positive beta coefficients (i.e., higher values in the ADHD or medication-exposed groups), and blue indicates negative coefficients. Subcortical regions not selected during the ML stage and therefore not included in LME modeling are coloured in grey.

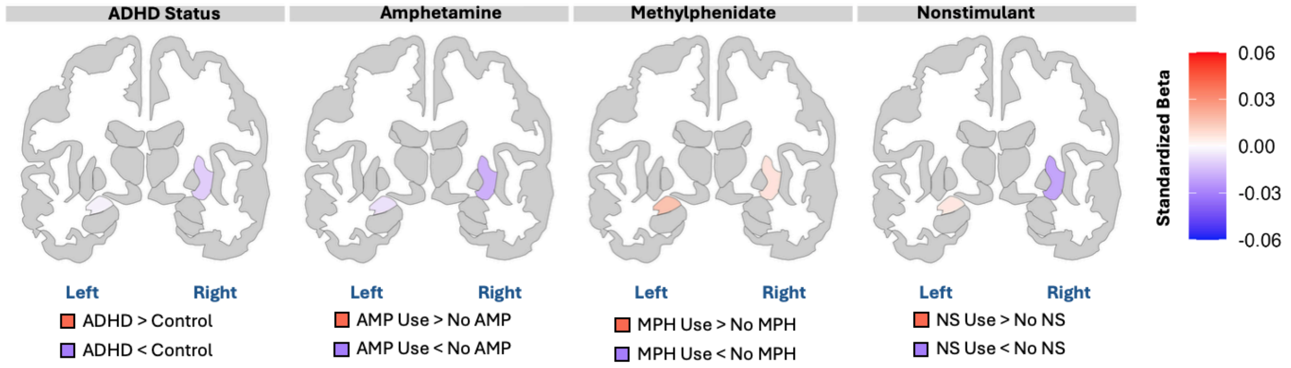


**Note:** The right accumbens area is not depicted in this section.
**Abbreviations:** AMP = amphetamine; MPH = methylphenidate; NS = nonstimulant.

**Supplemental Table 1.** Descriptive Statistics of the ABCD Study Population by ADHD Classification.

| | ADHD+RX[a] (N=1002) | ADHD-NoRX[b] (N=662) | Controls (N=10204) |
|---|---|---|---|
| **Age (years)** | | | |
| Mean (SD) | 119 (7.45) | 119 (7.49) | 119 (7.50) |
| Median [Min, Max] | 120 [107, 132] | 118 [107, 132] | 119 [107, 133] |
| Missing | 1 (0.1%) | 0 (0%) | 0 (0%) |
| **Sex at birth** | | | |
| Male | 732 (73.1%) | 444 (67.1%) | 5015 (49.1%) |
| Female | 270 (26.9%) | 218 (32.9%) | 5189 (50.9%) |
| **Race/Ethnicity** | | | |
| Non-Hispanic white | <550[c] | <340[c] | 5295 (51.9%) |
| Non-Hispanic Black | 170 (17.0%) | 117 (17.7%) | 1497 (14.7%) |
| Hispanic/Latinx | 158 (15.8%) | 128 (19.3%) | 2124 (20.8%) |
| Non-Hispanic Asian | <10[c] | <10[c] | 242 (2.4%) |
| Other | 124 (12.4%) | 79 (11.9%) | 1044 (10.2%) |
| Missing | 0 (0%) | 0 (0%) | 2 (0.0%) |
| **Household Income** | | | |
| <$50K USD | 340 (33.9%) | 204 (30.8%) | 2678 (26.2%) |
| $50K to $100K USD | 254 (25.3%) | 169 (25.5%) | 2645 (25.9%) |
| ≥$100K USD | 335 (33.4%) | 230 (34.7%) | 3996 (39.2%) |
| Don't know/Refuse to answer | 73 (7.3%) | 59 (8.9%) | 883 (8.7%) |
| Missing | 0 (0%) | 0 (0%) | 2 (0.0%) |
| **Highest Parent Education** | | | |
| < HS Diploma | <50[c] | <40[c] | 523 (5.1%) |
| HS Diploma/GED | 104 (10.4%) | 49 (7.4%) | 979 (9.6%) |
| Some College | 335 (33.4%) | 193 (29.2%) | 2546 (25.0%) |
| Bachelor Degree | 243 (24.3%) | 192 (29.0%) | 2578 (25.3%) |
| Post Graduate Degree | 281 (28.0%) | 196 (29.6%) | 3565 (34.9%) |
| Missing/Refused | <10[c] | <10[c] | 13 (0.1%) |
| **MRI Manufacturer** | | | |
| GE MEDICAL SYSTEMS | 213 (21.3%) | 144 (21.8%) | 2182 (21.4%) |
| Philips Medical Systems | 85 (8.5%) | 68 (10.3%) | 1220 (12.0%) |

| | | | |
|---|---|---|---|
| Siemens | 601 (60.0%) | 372 (56.2%) | 5902 (57.8%) |
| T1w Scan Not Usable or Incomplete | 103 (10.3%) | 78 (11.8%) | 900 (8.8%) |
| **Other Neuropsychiatric Meds** | | | |
| Yes | 163 (16.3%) | 32 (4.8%) | 108 (1.1%) |
| No | 839 (83.7%) | 630 (95.2%) | 10087 (98.9%) |
| Missing | 0 (0%) | 0 (0%) | 9 (0.1%) |

[a] ADHD+RX = medicated ADHD group (i.e., participants using at least one ADHD medication)
[b] ADHD-noRX = unmedicated ADHD group (i.e., participants who met ADHD diagnostic criteria and did not report ADHD medication use)
[c] Small cell sizes (< 10) were masked in accordance with ABCD data-use guidelines. Secondary cell suppression is implemented to ensure category frequencies cannot be recalculated by subtraction. Percentages for masked or suppressed cells are not shown.
**Abbreviations:** AMP = amphetamine; MPH = methylphenidate; NS = nonstimulant; HS = high school
